# An open-source, 3D-printable droplet microfluidics platform with strobe-enhanced microscopy

**DOI:** 10.64898/2026.10.19.758223

**Authors:** Pierre Padilla-Huamantinco, Matias Hurtado-Labarca, Christie Nel, Tomas Astudillo, Wladimir Araya, Juan José Leon, Josefina Belen Reyes Arroyo, Tobias Wenzel

## Abstract

Droplet microfluidics underpins many single-cell, precision biomedical, and nanomaterial techniques. However, implementing these methods reproducibly remains challenging outside specialised laboratories due to high equipment costs, operational inflexibility, mechanical impracticality at the microscope stage, as well as open-source pumping solutions that are inadequately validated for low-flow stability. We present an accessible, research-grade, open-source droplet microfluidics platform integrating a compact, microfluidics-optimised microscopy stage with networked syringe-pump control. Designed for local digital fabrication, primarily 3D printing, and widely available components, the platform supports reproducible builds across diverse settings. This setup enables stable droplet generation, observation of rapid droplet dynamics, and automation-ready networked workflows. The microscopy module uses a Raspberry Pi camera and microsecond stroboscopic illumination to capture sharp images of fast-moving droplets without the need for a high-speed camera. A modular hardware interface and extensible, web-accessible software stack provide a path to integrate additional modules and laboratory automation beyond the configuration evaluated here. For infusion, we introduce an open-source syringe pump optimised for low reliable flow rather than maximum rates, combining gearbox-assisted stepper mechanics with workflow-centric control that enables manual intervention (keypad control) and remote operation (web interface and automation hooks). We benchmark droplet regimes, size tuning, and time-stability under low-flow conditions (between 25 and 2500 µL/h) relevant to advanced assays and demonstrate biomedical readiness through the encapsulation and incubation of cells, beads, and bacteria in monodisperse droplets. The platform has been replicated across laboratories in Latin America and globally, including independent builds, supporting reproducibility. All design files, firmware/software, analysis notebooks, and benchmark datasets are openly released.

## Introduction

Droplet microfluidics is a cutting-edge technique that enables the generation and manipulation of customizable microenvironments across a range of sizes and throughputs for biological and chemical experiments. The technique facilitates rapid and parallelised assays while minimizing the reagent and sample volumes^1^. Among its biological applications, droplet microfluidics offers the scalable analysis of single cells, providing novel insights into individual cell behaviors, functional roles, and community-wide interactions in controlled environments^2,3^. Despite its potential to advance biomedical research, the widespread adoption of droplet microfluidics remains hindered by the reliance on costly proprietary instrumentation and complex experimental workflows^4^.

Most droplet research applications involve sample encapsulation into small monodisperse droplets (usually about 15 to 150 um diameter, pico- to nanoliter volume), requiring precise control of experimental parameters such as microchannel geometry and flow. Generating such microfluidic droplets requires at least two types of instruments: an imaging system with stroboscopic illumination^5^ or a high-speed camera^6^ that allows visualising the fast moving droplets, and a pumping system for an at least one aqueous and an oil phase^7^. Established pumping systems encompass syringe, peristaltic, and pressure pumps, where syringe pumps are widely regarded as the standard for laboratory-scale microfluidic research, because they are simple to operate, cost efficient compared to alternatives and provide less pulsation in their flow than peristaltic pumps.

Droplet microfluidic workstations usually rely on commercial proprietary instrumentation that requires a high up-front investment^7^, many with incompatible hardware and software that limits the automation and scalability of workflows across providers. Open-source (OS) instrumentation^8,9^ has emerged as an accessible and customizable alternative for imaging microfluidic operations and delivering various fluids and suspensions to microdevices. By employing open electronics^10^ (e.g., Arduino and Raspberry Pi) and digital manufacturing^11^ (e.g., 3D printing and laser cutting), peripheral instruments or integrated microfluidic workstations can be assembled within a few hours or days for a small fraction (typically 1-10%) of the cost. Nevertheless, reported OS instrumentation designs still exhibit limited functionality, performance, and validation for precision microfluidic applications such as single-cell assays, as well as the need for additional solutions to integrate and automate workstation modules.

In terms of imaging, OS designs typically implement a simple bright-field microscope based on a low-cost CMOS camera that allows the capture of the morphology and content of static or slow-moving microdroplets^12^. When droplets are produced at higher frequencies (e.g. >100 Hz) or move rapidly through microchannels, their motion blur increases considerably, hindering droplet monitoring and analysis^13^. Although the addition of stroboscopic illumination to imaging setups can resolve fast-moving particles^5,14,15^, there is no integrated OS implementation to our knowledge that combines a compact modular microfluidic workstation with a well-documented strobe-illumination. Compact instrumentation size benefits performance, time and cost savings through shorter single-use connection tubing and more efficient and flexible use of laboratory space.

OS syringe pumps are mainly 3D-printed, stepper motor-driven systems^9,16^. Available designs vary between stand-alone, dual-channel, and multichannel configurations. Reported microfluidic applications include the manipulation of single-phase fluids^17–19^, production of single and double emulsions^20–22^ including for microbial cultivation, and single-cell transcriptomics^23^. Unfortunately, the specific application data is often the only shared technical validation of the published instrument. Furthermore, OS syringe pumps often only report maximum flow rates and accuracy error^9^, leaving researchers with uncertainty about their suitability for the many applications that require a low but stable flow rate. In addition, response characteristics, such as the flow pulsation and stick-slip behaviour inherent to direct-drive stepper motors at low speeds, are rarely reported in a comparable manner^24^. From the many published options, we could not identify an OS pump design that met our flow stability and documentation expectations for our applications in droplets.

Proprietary pumps face a similar shortage of performance information, and standardised technical definitions and characterization workflows remain limited^9^, which makes model selection challenging and may give users an unjustified confidence to use some of these instruments for demanding applications. Industrial specifications in this area are still emerging^24,25^. Characterisation guides and further standardisation remains crucial for advancing hardware development and enabling reliable research^26^.

In this study, we present an accessible, research-grade, open-source droplet microfluidics platform that integrates a compact, microfluidics-optimised strobe-enhanced microscopy stage with networked syringe pump control. Designed for local digital fabrication, primarily 3D printing and laser cutting, and widely available components, this platform supports reproducible builds across diverse settings. Our syringe pumps were benchmarked against a premium proprietary pump to demonstrate comparable performance and reliability for droplet generation. In the absence of clear characterisation guidelines and equipment standards in the field, the platform was benchmarked under different microfluidic conditions for droplet generation, particle encapsulation, incubation, and automated dilutions, demonstrating proof of principle images, strobe performance, droplet monodispersity, droplet content analysis, and flow sensor data samples to illustrate flow stability.

## Platform design and fabrication

We present a droplet microfluidics platform comprising two instruments: a strobe-enhanced microscopy stage for fast imaging of droplets and 3D-printed syringe pumps for multiphase fluid manipulation, see **Fig. 1A,B**. Both instruments can be controlled from a joined web-based graphical user interface (**Fig. 1C**) or via an API. The syringe pump additionally features a manual keypad interface for fast manual adjustment and intervention in experiments. The microscopy stage is based on a custom electronics board named “Rio-controller” (for Spanish “River”) that plugs onto a Raspberry Pi 4 single-board computer. The board supports several microcontroller modules including strobe control, and additional modules to interface with pressure controllers, flow sensors and heaters (**Fig. 1D left**). An earlier configuration of these electronics and software was combined with pressure-driven flow, flow sensing, and sample and chip heating, to support a gel-microdroplet assay of bacterial plasmid stability^27^. This application illustrates the reconfigurability of the controller architecture. The syringe pump is based on a separate custom board with a ESP32 microcontroller that, in addition to the manual keypad, can be controlled by wifi and USB (**Fig. 1D right**). The platform modularity is supported by a highly modular software design, hardware simulation options for testing, and detailed documentation for software and hardware, including a step-by-step assembly guide suitable for users without technical experience (**Fig. 1E**).

**Figure 1.**
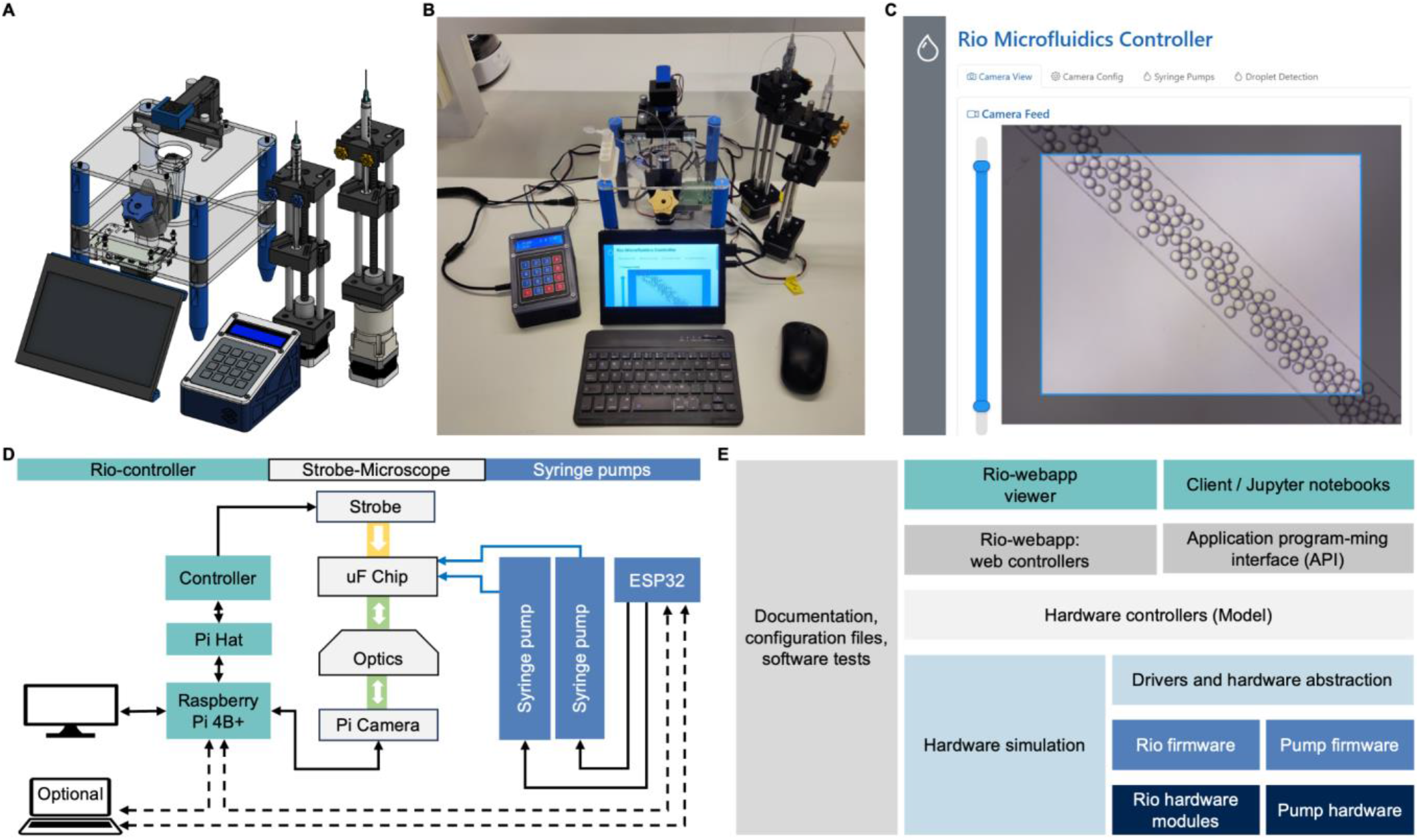
Architecture of the open droplet microfluidics workstation. **(A)** CAD design and rendering of the modular system including the strobe microscope stage, controller electronics, and syringe pumps. **(B)** Photograph of the assembled workstation during operation. **(C)** Screenshot of the web-based graphical user interface used for device control and monitoring. **(D)** Hardware communication diagram showing the Raspberry Pi–based “Rio-controller” interfacing with the strobe microscopy stage and the syringe pumps driven by an ESP32 controller. **(E)** Overview of the modular software architecture including hardware modules and firmware, drivers, hardware abstraction and simulation, controller layer, web application and API interfaces, and supporting documentation and configuration files.

The microscopy stage was designed to permit visual assessment of rapid droplet formation using low-cost camera hardware. Conventional systems typically freeze droplet motion using high-speed cameras or, less commonly, stroboscopic illumination integrated into commercial microscopes^5,14,28,29^, at high instrument cost and complexity. Increasing the camera frame rate and shortening its electronic exposure can reduce motion blur, but also decreases photon collection and may restrict image size or signal-to-noise ratio^30^, particularly when using sensors with small pixels <5 µm. Here we combined inexpensive cameras with synchronised microsecond LED illumination in our minimal, fluidics-friendly microscopy stage. When the illumination pulse occurs within the sensor integration window, the pulse width determines the effective optical exposure. Sharp snapshots of rapidly moving droplets can therefore be acquired without high-speed image acquisition.

The imaging stage is a derivative of the OpenFlexure microscope project^31^, specifically the flat-top stage (https://rwb27.gitlab.io/openflexure-flat-top-microscope/). It is designed to be compact with a clean, flat top that supports microfluidic chips of all sizes, is robust to applied pressure from tubing exchanges and other typical interventions, and is easy to clean from spillages with no cracks or gaps between mechanisms that are typical of other microscopy stages. The OpenFlexure 3D printed mechanism^31^ allows fine adjustment of the camera focus that allows a Z-axis travel range of 4 mm and a step size as low as 50 nm and is compatible with optical configurations developed for other OpenFlexure projects, such as objectives, webcams, and fluorescence mini-cubes, as far as the space inside the stage allows. However, here the optics is kept simple for inverted transmission wide-field microscopy that allows to find small microfluidic channels with ease within the large area of chips, instead of large magnifications high quality optics to e.g. characterise individual cells. The sensor used here is a Raspberry Pi Camera V2 (Sony IMX219, 8-megapixel CMOS sensor), coupled with a 3D-printed tube lens holder and a standard 4×/0.10 NA objective, offering a 1.4 x 1 mm field of view (FOV) with a 5 mm working distance. In our view and experience, it is more effective to separate droplet generation and quantitative imaging set-ups to optimise for their respective tasks. We demonstrate simple quantification tasks that can be performed in the process of droplet generation, but more precise quantification and advanced tasks would require changes to the optics that would render droplet generation less convenient and increase cost.

The stage controller was implemented using a Raspberry Pi 4B board, a custom-designed hardware-attached-on-top (HAT) board, and a custom PIC-based (PIC16F18856) strobe controller. The stroboscopic illumination comprised a 1W neutral white LED coupled with a polymer condenser lens. In this strobe-centered configuration, the pulse duration controls the effective exposure time of the sensor at the available sampling rates.

The syringe pump system is a derivative of a stepper motor-based open-source (OS) pump design by Samokhin et al.^19^, which has similar technical features to the majority of other 3D printed syringe pump designs and was deemed aesthetically pleasing. We re-designed the missing CAD files, optimised them with heat inserted threads, and developed a new controller module plus a gear-box version of the pumps. Two syringe pump versions were implemented: a low-cost syringe pump close to the original design (direct drive), and a syringe pump with a 100:1 planetary gearbox (EG17-G100, StepperOnline) to increase effective torque and angular resolution (**Fig. 1A**).

The pump controller is based on a custom PCB (printed circuit board), breadboard alternative available, with an ESP32 microcontroller and Trinamic (TMC-series) motor drivers, within a compact footprint. The physical interface was implemented using an LCD display and a 4 × 4 keypad. The controller supports up to four pumps for multiphase control.

Local digital manufacturing, such as 3D printing and laser cutting, for producing most mechanical structures, and widely available electronic components for sensing, actuation, and control, were prioritised to ensure accessibility and customization without relying on specialised supply chains. The carriage and motor mounts, as well as the controller cases for both pump models, have successfully been fabricated by 3D printing with PLA (polylactic acid) or more temperature resistant PETG (polyethylene terephthalate glycol) filament.

Modularity was incorporated at hardware and software levels to use stage and pump from a single interface or separately in different research and education settings. The capabilities of the instruments can be modified or upgraded gradually by users depending on local requirements or available resources; for example upgrading from a Raspberry Pi camera lens to a microscopy objective, or operating either one direct-drive pump or multiple gearbox-assisted pumps using the same pump controller. This flexible and incremental development approach lowers the initial barrier to adoption while allowing for future hardware extensions. At the software level, instrument control is provided through a web-based interface and API (application programming interfaces), allowing operation across multiple operating systems without dependence on platform-specific software.

## Results and discussion

### Stroboscopic droplet imaging with low-cost hardware

To evaluate the capability of the platform to visualise droplet production, identify of flow regimes, and estimate of droplet size and spacing, we compared conventional illumination and synchronised stroboscopic imaging using the Raspberry Pi Camera V2 of the platform (**Fig. 1**), with an entry-level industrial high-speed camera (MAKO-U029B, Allied Vision). Both cameras were equipped with the same 4× 0.10 NA objective and equivalent optical paths. Droplets were generated at approximately 9 kHz for imaging, using an oil-to-aqueous flow-rate ratio of 2000:1000 μL h^−1^ (**Fig. 2**).

**Figure 2.**
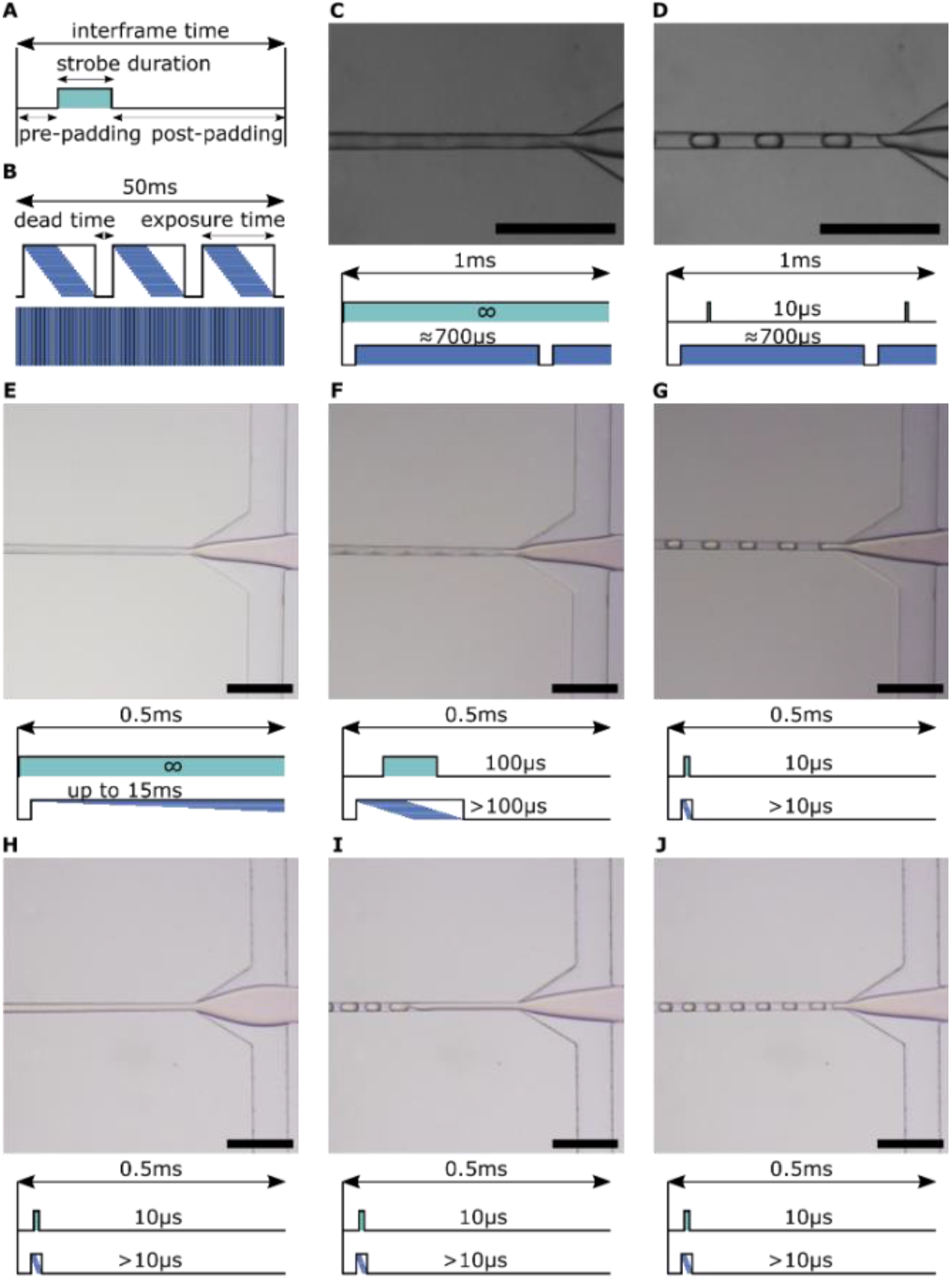
Strobe illumination imaging and exposure schematics. **(A)** Timing diagram of a strobe pulse within one frame interval showing pre-padding, strobe duration, and post-padding. **(B)** Exposure mechanisms of the Raspberry Pi Camera V2 at 60 FPS with rolling shutter (line-by-line readout) and the MAKO-U029B industrial camera at 1359 FPS with global shutter. **(C**,**D)** MAKO camera images with **(C)** continuous illumination and **(D**) 10 µs strobe illumination synchronised to the camera trigger. **(E–J)** Images acquired with the Raspberry Pi Camera V2 under different strobe and flow conditions. Schematics below each image indicate strobe timing and estimated auto-adjusted exposure times. **(E)** Continuous illumination with long exposure. **(F)** 100 µs strobe illumination. **(G)** 10 µs strobe illumination. **(H–J)** 10 µs strobe illumination at different oil-to-aqueous flow ratios showing **(H)** tubing, **(I)** jetting, and **(J)** dripping flow regimes. All scale bars: 200 µm.

Under continuous illumination (no strobe), the industrial camera operating at high frame rates above 1000 fps (exposure < 1 ms) was unable to resolve droplets clearly (**Fig. 2C**). The same limitations of course also applied to the slower Raspberry Pi Camera V2 operating at 60 fps (frames per second) with an exposure time up to 5 ms (frame interval: ∼17 ms) (**Fig. 2E**).

Synchronised microsecond stroboscopic illumination decouples motion freezing from the camera frame rate and nominal exposure time, see schematic in (**Fig. 2A, B**). Single high-intensity illumination pulses (∼10 µs) enabled clear visualization of droplet formation with both cameras (**Fig. 2C–G**). The industrial camera achieved higher temporal resolution but required operation at a reduced image size (320 × 240 pixels) for high frame rates. The low-cost Raspberry Pi Camera V2 acquired qualitatively comparable full images at 1024 × 768 pixels at 60 fps. While both sensors feature small sensors (1.12 µm pixel CMOS chip for the Raspberry Pi Camera V2), the Raspberry Pi camera additionally features a rolling shutter mode (schematic in **Fig. 2B**). This complicates strobe based image capture because each pixel row is read out sequentially. If the strobe pulse is significantly shorter than the exposure time, only some rows receive light during the readout, leaving parts of the image dark. However, thanks to the automatically adjusted exposure times of the camera down to 9 µs, all images appeared fully illuminated and successfully resolved droplets generated at frequencies exceeding 9 kHz. These findings are consistent with previous reports demonstrating that global shutters are not inherently required for stroboscopic imaging^15^. The presented synchronised stroboscopic illumination enables low-cost sensors to perform blur-free imaging tasks traditionally requiring expensive ultra high-speed cameras with acquisition rates up to 100,000 fps.

To evaluate the motion blur effect with the Raspberry Pi Camera further, images were acquired using strobe pulse widths of 100 µs and 10 µs. Motion blur can be approximated as the product of droplet velocity and effective exposure time. The average droplet velocity was estimated to be 1 m s^−1^ (flow 3000 μL h^−1^, channel width ∼24 µm , height ∼35 µm). Consequently, a 100 µs illumination pulse corresponds to a motion blur of approximately 100 µm, comparable to the measured inter-droplet spacing of 110 µm. Under these conditions, droplets were highly blurred, preventing reliable identification of individual droplets (**Fig. 2F**). Reducing the pulse duration to 10 µs decreased the theoretical motion blur to 10 µm, and sharp droplet boundaries were consistently resolved at this high flow rate even at the flow-junction, the narrowest section of the chip (**Fig. 2G**). Note that droplet quantification was performed at broader sections of the chip where the droplet speed is 10x reduced.

The improved image quality obtained with short stroboscopic pulses also enabled direct visualization of distinct flow regimes (**Fig. 2H–J**) and qualitative assessment of droplet size, morphology, and inter-droplet spacing under different flow conditions (**Fig. 2G, J**). These observations demonstrate that synchronised stroboscopic illumination provides sufficient image quality not only for visualizing fast-moving droplets but also for supporting routine experimental tasks such as monitoring droplet stability and identifying operating conditions during microfluidic experiments.

### Research-grade open-source syringe pumps

The accuracy and quality of fluid delivery from syringe pumps is determined by multiple factors, including mechanical transmission, torque availability vs. backpressure, backlash, compliance, and stepper motor induced pulsations. To evaluate the quality of the proposed open-source (OS) syringe pump for advanced droplet research applications, we characterised (**Fig. 3**) its volumetric flow accuracy, temporal stability, and frequency-domain dynamics. We benchmarked the gearbox assistant pump (**Fig. 3I**) performance against our standard OS direct-drive design (**Fig. 3F**) and a premium proprietary syringe pump (Harvard Apparatus PHD2000, **Fig. 3C**). Flow measurements were performed with a precision flow sensor (Sensirion LG16-0431D with 5% accuracy between 120 and 4800 μL h^−1^) upstream of the droplet generation chip .

**Figure 3.**
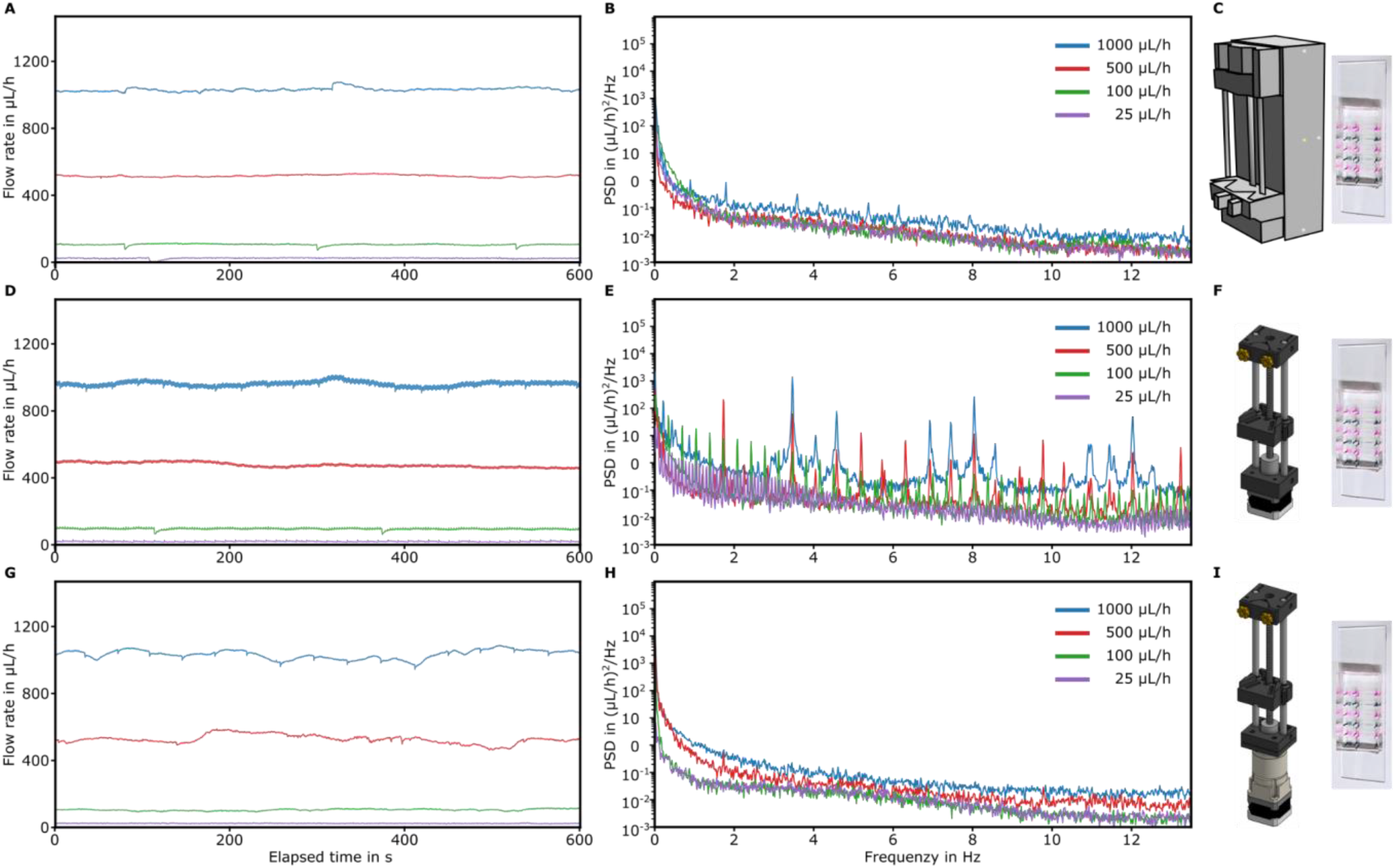
Flow rate stability and spectral analysis for a high-resistance microfluidic chip across pump configurations and target flow rates at high resistance. Time series (left column) and power spectral density (PSD, right column) of flow rate measurements at four target flow rates (1000, 500, 100, and 25 µL h^−1^) for three syringe pump configurations: (A) Harvard Apparatus reference pump, (B) OS Pump without gearbox, and (C) OS Pump with gearbox. Time series show 600 s of continuous acquisition at steady state. PSD plots were computed from mean-subtracted flow signals via Welch’s method.

All three pump platforms exhibited a linear relationship between the target and measured flow rates across the tested operating range (**Fig. 3A, D, G**), with coefficients of determination R^2^ exceeding 0.9999. The direct-drive pump maintained a mean error of 8%, whereas both the commercial and gearbox-assisted pumps slightly exceeded the target flow rates, with mean errors below 3%. These results suggest that the increased torque provided by the 100:1 planetary gearbox improves the accuracy of fluid delivery, yielding performance comparable to that of the commercial reference pump under all four experimental conditions. Despite small offsets, both OS designs remained close to the accuracy of the flow sensor and within the flow accuracy range typically reported for commercial syringe pumps^32^.

While flow accuracy and coefficient of variation provide useful performance metrics, they do not reveal the mechanical origin of flow fluctuations. To investigate these dynamics, the measured flow signals were analysed using Welch power spectral density (PSD) (**Fig. 3C, F, I**). The PSD was computed using a Hann window with 50% overlap, providing a frequency resolution of Δf = 0.0132 Hz (segment duration ≈ 76 s), sufficient to resolve stepping frequencies for low flow rates, while separating slower drift processes associated with friction events, plunger creep, and thermal expansion.

The premium proprietary pump delivered comparatively smooth flow profiles; nevertheless, discrete steps for lowest flow rates and transient disturbances remained evident (**Fig. 3A, B**). Our highest-performing, gearbox-driven configuration achieved similarly stable delivery (**Fig. 3G, H**). Both architectures maintain high plunger force while increasing the number of motor steps per unit displacement: the commercial pump combines a toothed timing-belt reduction drive with a specialised fine-pitch lead screw, whereas our design obtains a greater mechanical reduction through a 100:1 gearbox coupled to a standard lead screw. The downstream hydraulic resistance imposed by small-cross-section microfluidic channels further attenuated the residual pulsations, rendering their effect on delivered flow negligible (**Fig. 3B, G**). These results show that step-related flow disturbances are a general feature of stepper-driven syringe pumps rather than a limitation unique to OS designs.

The direct-drive pump exhibited pronounced stepping harmonics whose frequencies increased proportionally with flow rate, consistent with a kinematic origin. Dominant peaks were observed at 1.73 Hz (PSD power = 209.73 (μL h^−1^)^2^ Hz^−1^) and 3.47 Hz (PSD power = 1,417.40 (μL h^−1^)^2^ Hz^−1^) for flow rates of 500 and 1000 μL h^−1^, respectively (**Fig. 3D, E**). The integration of a 100:1 planetary gearbox significantly altered the dynamic behaviour of the pump (**Fig. 3G, H**). By reducing the linear step size from 313 to 3 nm, the theoretical stepping frequency increased by two orders of magnitude (for example, from 7.55 Hz to approximately 755 Hz at 500 μL h^−1^), shifting the dominant mechanical excitation toward substantially higher frequencies. Rather than eliminating stepping-induced pulsations, the gearbox redistributes their spectral energy toward higher frequencies that are effectively filtered by the hydraulic system. In short, the addition of a gearbox^18^ to the direct drive design typical for open source pump designs^9^ delivers a much smoother flow with significantly reduced flow pulsation within a large range of flow rates tested here (**Fig. 3E** vs **H**). These architecture-specific spectral patterns were retained across resolutions of 8, 32, and 256 microsteps (**Fig. S1**).

The present analysis demonstrates that a commercial syringe pump also exhibits measurable stepping harmonics, highlighting the importance of characterizing syringe pump dynamics using frequency-domain methods rather than relying exclusively on steady-state flow measurements, as reported in previous studies for this pump model^33,34^. Pulsation attenuation should also be re-validated under microfluidic systems with different hydrodynamic resistances, where the filtering characteristics are expected to vary^35^. The presence of pump-induced pulsations may become particularly relevant in applications with limited hydraulic damping, such as organ-on-chip perfusion systems, or in biological assays involving shear-sensitive cells, where periodic flow oscillations may directly influence cellular responses^36^.

### Droplet generation and monodispersity

Droplet generation is the primary application of the proposed microfluidic platform, which requires stable flow to generate monodisperse droplets across a broad range of operating conditions. Because droplet uniformity can be highly sensitive to fluctuations in flow delivery, we evaluated whether the improved flow characteristics of the gearbox-assisted syringe pump translated into enhanced monodispersity of droplets. Droplet diameter distributions were quantified using the droplet generation chip and stroboscopic imaging at four oil-to-aqueous flow rate conditions. For the experiment, a constant 2:1 oil to water flow ratio was maintained, while the absolute flow rate was changed over approximately one and a half orders of magnitude (**Fig. 4**).

**Figure 4.**
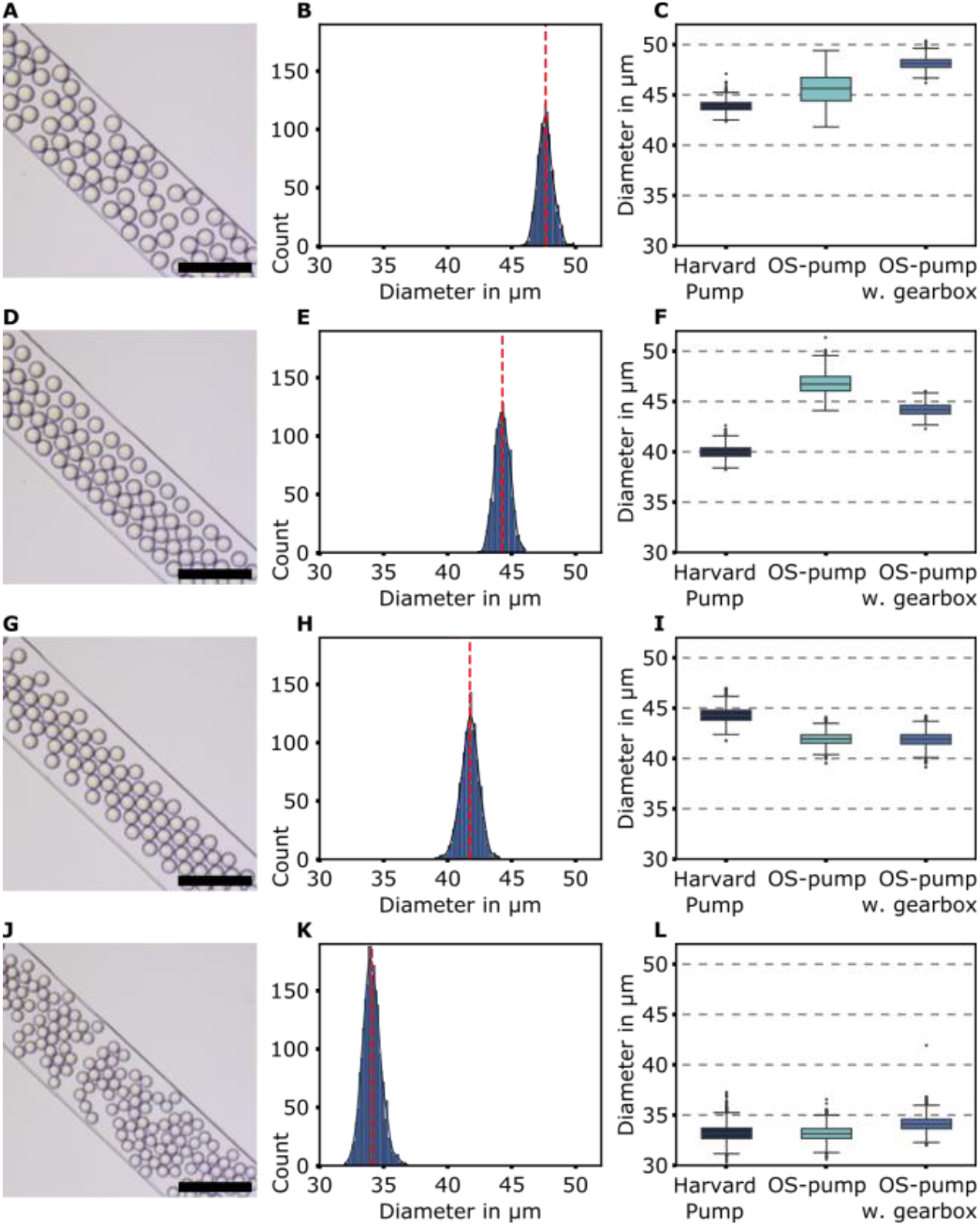
Droplet diameter distributions across flow rates and pump types at a fixed 2:1 oil-to-aqueous flow ratio. **(A,D,G,J)** Representative images of droplet generation using the open-source syringe pump with gearbox. Scale bars: 200 µm. **(B,E,H,K)** Corresponding droplet diameter histograms; red dashed lines indicate mean values. A total between 1121 and 2172 droplets were analysed per condition (10 images each). **(C,F,I,L)** Box plots of droplet diameters showing interquartile range. Data from the gearbox pump are compared to a proprietary Harvard Apparatus pump and an open-source pump without gearbox. Aqueous flow rates were **(A–C)** 25 µL h^−1^, **(D–F)** 100 µL h^−1^, **(G–I)** 500 µL h^−1^, and **(J–L)** 1000 µL h^−1^, with oil flow rates set to maintain a 2:1 ratio.

Representative bright-field images obtained with the gearbox-assisted pump and stroboscopic imaging system demonstrated stable dripping behaviour under all operating conditions (**Fig. 4A, D, G, J**). As expected for flow-focusing geometries, increasing the absolute flow rate reduced droplet diameter owing to the corresponding increase in capillary number^35^. Across all conditions, droplet size distributions remained narrow (**Fig. 4B, E, H, K**), with within-replicate coefficients of variation below 3%, matching the monodispersity typically required for droplet microfluidic applications^37^. The droplet generation frequencies were approximately 229 Hz, 918 Hz, 4.6 kHz, and 9.2 kHz for oil-to-aqueous flow rates of 50:25, 200:100, 1000:500, and 2000:1000 μL h^−1^, respectively. In short, the platform supported stable high-throughput droplet production within a wide range of flow rates.

To determine whether pump architecture influenced droplet production, the gearbox-assisted pump was again compared to the direct-drive configuration representative of widely reported 3D-printed syringe pumps^9^ and the proprietary Harvard Apparatus PHD2000 syringe pump. At the two highest flow-rate conditions (1000:500 and 2000:1000 μL h^−1^), all three platforms produced nearly identical droplet diameters and high monodispersity (**Fig. 4I, L**), with coefficients of variation ranging from 1.4% to 3.2%. These results indicate that, under operating conditions where hydrodynamic damping effectively suppresses residual pump pulsations, both OS designs achieve droplet quality comparable to the premium proprietary instrument. However, differences between OS pump architectures became evident under lower flow-rate conditions. At 200:100 μL h^−1^ (**Fig. 4F**), the gearbox-assisted pump achieved substantially lower variability (CV = 2.0%) than the direct-drive configuration (5.2%). At 50:25 μL h^−1^ (**Fig. 4C**), the trend persists (CV of 1.4% vs 2.5%). The premium proprietary pump behaved similarly to the gearbox-assisted pump model. Here we only compare droplet polydispersity and not average droplet sizes, as experiments were conducted on different days using different microfluidic channels and chips. Potential contributors to size differences include manual connections to chips, variations in room temperature, or chip-to-chip geometric variability. However, replica experiments (**Fig. S2**) confirm monodispersity trends.

The observed droplet-generation behaviour is consistent with the flow characterization presented in the previous subsection. The pulsations measured in the direct-drive pump at low flow rates appear sufficient to increase droplet-size variability. In contrast, the gearbox-assisted design preserved the low variability expected from its substantially reduced pulsation spectrum, indicating that smoother mechanical actuation directly improves droplet uniformity at least at lower flow rates. The results are consistent with those of Booeshaghi *et al*. (18), who reported variability between the direct-drive OS Poseidon syringe pump and a Harvard Apparatus pump. Our results demonstrate that mechanically optimised OS syringe pumps can serve as a practical alternative for premium proprietary hardware in advanced droplet-generation experiments.

### Automated experimental workflows

Beyond accurate fluid delivery, our droplet microfluidics platform design supports dynamic control and programmable experimental workflows while preserving modularity. To demonstrate the automation capability of the platform, the gearbox-assisted syringe pumps were operated in a synchronised three-pump configuration together with the microscopy system through a Jupyter notebook script using the platform’s API interface. Two complementary automated workflows were evaluated: (i) programmable droplet-size modulation by varying the continuous-phase flow rate (**Fig. 5**), and (ii) concentration modulation by varying the relative flow rates of two aqueous streams while maintaining a constant total dispersed-phase flow rate (**Fig. 6**).

**Figure 5.**
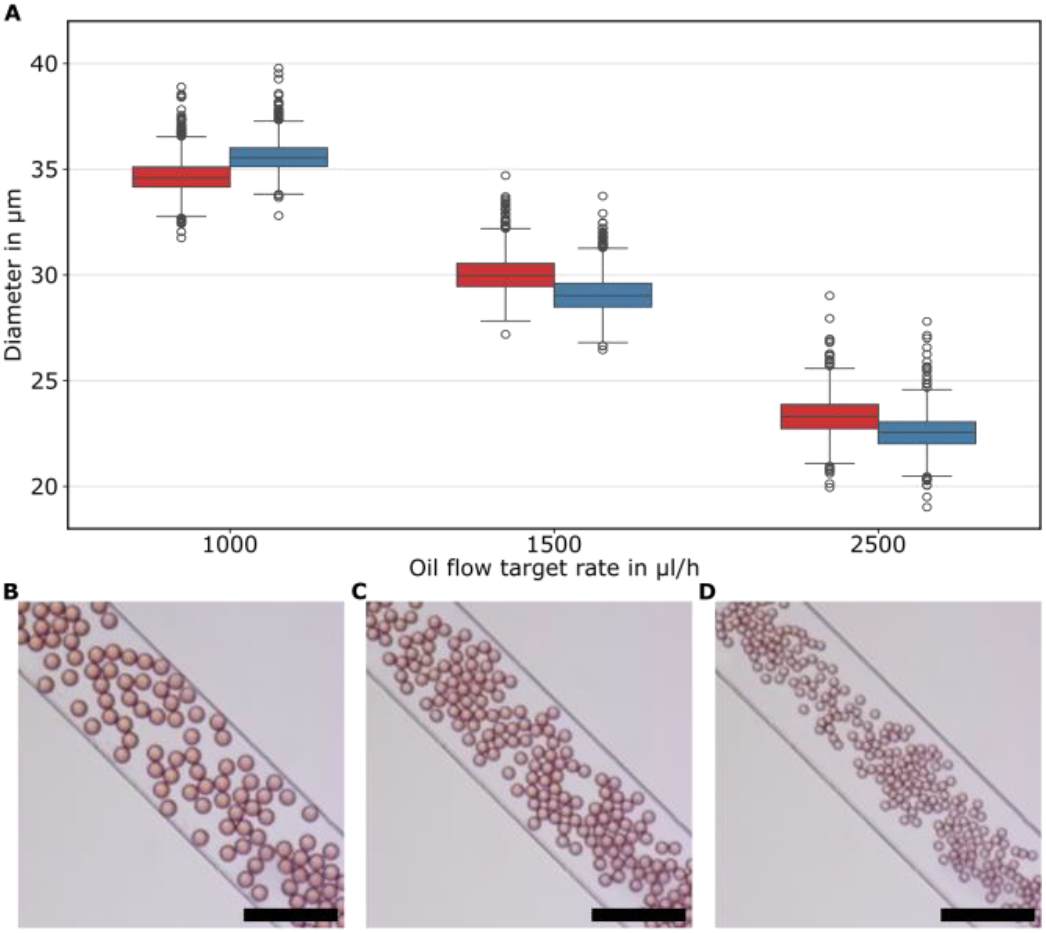
Droplet size modulation using three high-torque OS pumps. Representative images of automated droplet size modulation using the open-source syringe pump with gearbox. Scale bars: 200 µm. (A) Box plots of droplet size distribution at different oil flow rate targets. Forward (increasing oil flow rate) and backward (decreasing oil flow rate) sequences were performed to compare reproducibility, Oil flow rates were (B) 1000 µL h^−1^, (C) 1500 µL h^−1^, and (D) 2500 µL h^−1^, with total aqueous flow rates of 500 µL h^−1^ (Q_water_ = 250 μL h^−1^ and Q_dye_ = 250 μL h^−1^).

**Figure 6.**
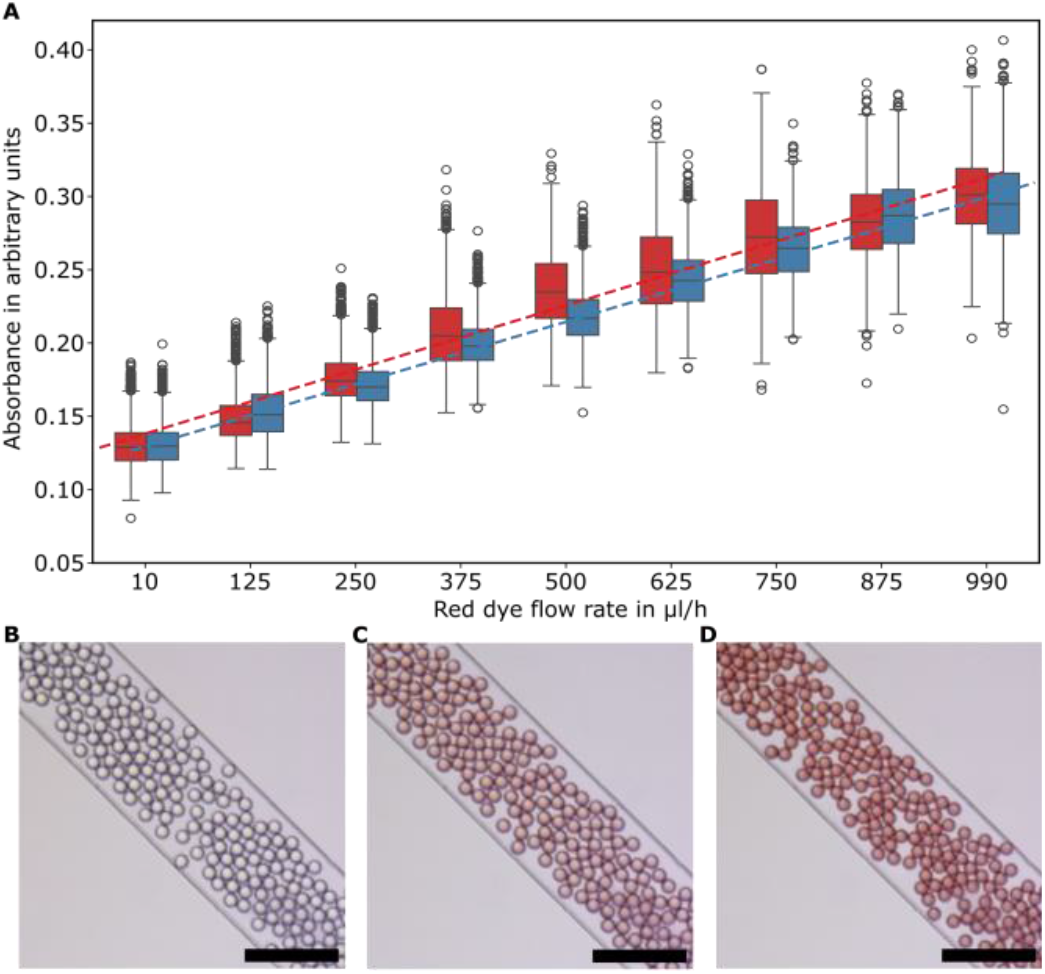
Automated dilution modulation using three high-torque OS syringe pumps. (A) Box plots of droplet absorbance as a function of red dye flow rate (10–990 µL h^−1^), with the total aqueous flow rate maintained at 1000 µL h^−1^ and the oil flow rate fixed at 1500 µL h^−1^. Red and blue box plots correspond to the Forward (increasing flow rate) and Backward (decreasing flow rate) sequences, respectively, demonstrating close agreement between both trajectories and confirming reversible and reproducible dilution control. (B–D) Representative brightfield micrographs of droplets at three dye compositions: low (Q_dye_ = 10 µL h^−1^), intermediate (Q_dye_ = 500 µL h^−1^), and high (Q_dye_ = 990 µL h^−1^) red dye fractions, with continuous phase flow rate Q_oil_ = 1500 µL h^−1^ and complementary aqueous flow Q_water_ maintained at 990, 500, and 10 µL h^−1^, respectively. Scale bars: 200 µm.

Droplet size was modulated by automatically stepping the oil-phase flow rate through three programmed values while maintaining constant aqueous flow rates (**Fig. 5**). Representative bright-field images acquired at each operating point (**Fig. 5B-D**) show stable droplet generation with uniformly spaced droplets whose diameter decreased progressively as the oil flow rate increased, consistent with expectations. Automated image analysis of 9,486 droplets confirmed this behaviour quantitatively (**Fig. 5A**). Mean droplet diameter decreased from 35.1 μm at 1000 μL h^−1^ (n = 3414) to 29.5 μm at 1500 μL h^−1^ (n = 4101) and 22.9 μm at 2500 μL h^−1^ (n = 1971), corresponding to an overall reduction of approximately 35% across the tested operating range. Despite these dynamic flow transitions, droplet monodispersity remained high, with coefficients of variation of 2.2%, 2.9%, and 4.1%, respectively, remaining below the 5% threshold for droplet monodispersity.

The reversibility of the automated control was evaluated by comparing increasing (forward) and decreasing (backward) flow-rate sequences. Both trajectories produced nearly identical droplet-size distributions (**Fig. 5A**), with mean diameters differing by less than 3% across all operating points. At oil flow rates of 1000, 1500, and 2500 μL h^−1^, forward and backward sequences yielded closely matched mean diameters of 34.7 versus 35.6 μm, 30.0 versus 29.1 μm, and 23.4 versus 22.6 μm, respectively. This lack of hysteresis indicates that the gearbox-assisted actuation responds reproducibly to changes in commanded flow rate.

To demonstrate automated control of droplet composition independently of droplet size, the oil flow rate was fixed at 1500 μL h^−1^ while maintaining a constant total dispersed-phase flow rate of 1000 μL h^−1^ (**Fig. 6**). The relative flow rates of water and red dye were then automatically programmed through nine sequentially conditions ranging from 990:10 to 10:990 μL h^−1^. Maintaining a constant total dispersed-phase flow rate ensured that changes in optical intensity reflected only variations in dye concentration rather than changes in droplet volume. Representative images show the resulting concentration gradient, with droplets transitioning from nearly transparent to deeply coloured as the dye fraction increased (**Fig. 6B-D**). Quantitative image analysis showed a steady decrease in green-channel intensity from approximately 140 to 95 arbitrary units as the dye flow rate increased, reflecting the complementary absorption of the red dye. Its proportionality was confirmed by converting absorption into absorbance using the Beer–Lambert relationship (**Fig. 6A**), providing an independent quantitative representation of the programmed concentration gradient. For reference, the absorbance distributions of manually stablished dye-to-water ratios were consistent with the results of the automated assay (**Fig. S3**).

Forward and backward concentration sweeps (**Fig. 6A**) produced nearly identical responses in both green-channel intensity and calculated absorbance, with closely matching medians and interquartile ranges across all nine programmed conditions, again showing negligible hysteresis.

These results demonstrate programmable microfluidic workflows integrating both modular OS instruments through coordinated multi-pump control and imaging without manual intervention. The automated continuous reconfiguration of droplet microenvironments during a single experiment is directly applicable to dose–response studies, reagent-gradient generation, combinatorial encapsulation, and other automated single-cell workflows, providing the foundation for advanced high-resolution biological experiments.

### Proof-of-principle biological applications

To demonstrate that the proposed platform supports biologically relevant microfluidic workflows for the encapsulation of more challenging buffers, particles and cells, we performed a series of representative proof-of-principle applications using the gearbox-assisted syringe pump and the strobe-enhanced microscopy stage (**Fig. 7**).

**Figure 7.**
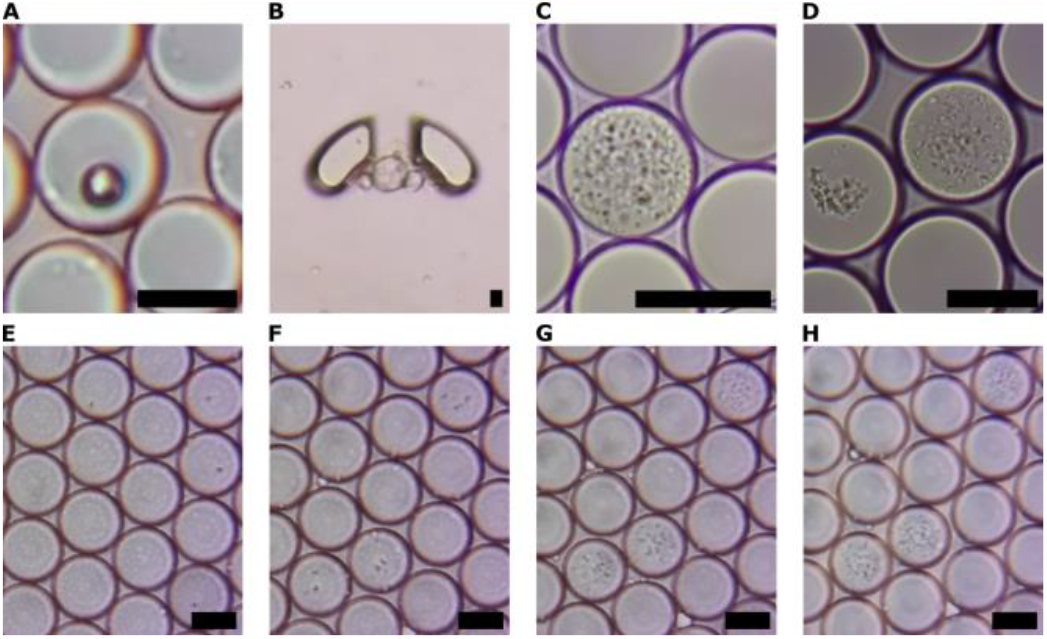
Biological validation of the open droplet microfluidics workstation using on-stage microscopy. **(A)** Encapsulation of individual polystyrene beads. **(B)** Flow-trapping of a multicellular human organoid in a microfluidic chip. **(C,D)** Bacterial growth originating from single-cell encapsulation, showing *Staphylococcus aureus* **(C)** and *Escherichia coli* **(D)**, with distinct intra-droplet growth behaviors (dispersion vs. aggregation). **(E–H)** Time-resolved growth of *S. aureus* following single-cell encapsulation at 0 h **(E)**, 4 h **(F)**, 8 h **(G)**, and 12 h **(H)**, demonstrating sustained clonal expansion within droplets during on-stage incubation at room temperature. All images were acquired using the presented workstation. Scale bars: 25 µm.

Encapsulation of individual polystyrene beads was first performed as a controlled benchmark for single-object loading into droplets (**Fig. 7A**). The bright-field image shows a single encapsulated 10.5 μm bead. Although biologically inert, microbeads are widely employed in droplet microfluidics as cell surrogates for validating flow-focusing designs^38^ barcoding and imaging correlation^39,40^, or sensing applications^41^.

To demonstrate compatibility with alternative microfluidic workflows, a multicellular human gastric organoid was captured in a reversible microfluidic trapping device (**Fig. 7B**). Successful immobilization at low flow rates together with clear visualization of the organoid illustrates that the platform is not restricted to droplet-generation workflows but can also support trapping-based microfluidic devices for the handling and imaging of three-dimensional biological structures and compatibility with perfusion cell culture.

The platform was next evaluated in a representative single-cell microbiology workflow by encapsulating two bacterial model organisms exhibiting distinct in-droplet growth phenotypes. *Staphylococcus aureus* formed compact microcolonies that progressively occupied the droplet interior as dispersed aggregates (**Fig. 7C**), whereas sfGFP *Escherichia coli* developed more tightly clustered colonies with a clearly different spatial organization (**Fig. 7D**). The ability to distinguish these contrasting growth morphologies directly from bright-field images demonstrates that the custom microscopy stage provides sufficient spatial resolution and image contrast for simple phenotypic characterization at the single-droplet level, even if we often use specialised setups for it.

The bacterial growth of *S. aureus* was monitored over a 12 h in-droplet incubation period using time-lapse bright-field imaging (**Fig. 7E–H**). Droplets containing individual encapsulated cells were readily identified (**Fig. 7E**). After 4 h, early microcolony formation became apparent (**Fig. 7F**), followed by continued proliferation after 8 h (**Fig. 7G**). By 12 h, mature microcolonies occupied a substantial fraction of the droplet volume (**Fig. 7H**). Throughout the incubation period, the droplet monolayer remained stable, with no visible evidence of evaporation or droplet coalescence, indicating that the sealing protocol maintained stable incubation conditions during extended room-temperature observations. The microscopy stage consistently resolved bacterial cells within increasingly crowded droplet interiors using bright-field imaging alone. Such long-term monitoring of encapsulated microorganisms is a fundamental requirement for droplet-based assays such as microbial growth studies, antibiotic susceptibility testing, and functional single-cell analyses.

Collectively, these proof-of-principle applications demonstrate that the proposed platform supports a diverse range of biologically relevant microfluidic workflows extending beyond droplet production alone. The successful encapsulation of inert particles, manipulation of multicellular organoids, and monitoring of bacterial growth illustrate that the integrated pumping, imaging, and control systems can support applications requiring precise fluid handling, stable droplet generation, automation, and prolonged simple microscopic observation.

### Replicability across laboratories

This open-source droplet microfluidics platform was designed with replicability as a core objective, alongside improving accessibility and customization for microfluidic research. To achieve this, open-source design principles and recommendations^8,11,20^ were applied to the platform architecture. Consistent with the OSHWA definition and best practices^42^, source files, BOM (bill of materials), and documentation were publicly released for community replicability. All design files are provided in both native editable and derived fabrication-ready formats such as STL-files, facilitating the rebuilding as well as the modification of the platform for different use cases (**Fig. 8A-D**). A complete BOM accompanies each instrument, listing quantities, suppliers, and part numbers when available. Comprehensive assembly instructions were implemented using the hardware documentation tool GitBuilding^43^ on GitHub. All hardware resources are released under the CERN Open Hardware Licence v2, Weakly Reciprocal (CERN-OHL-W-2.0).

**Figure 8.**
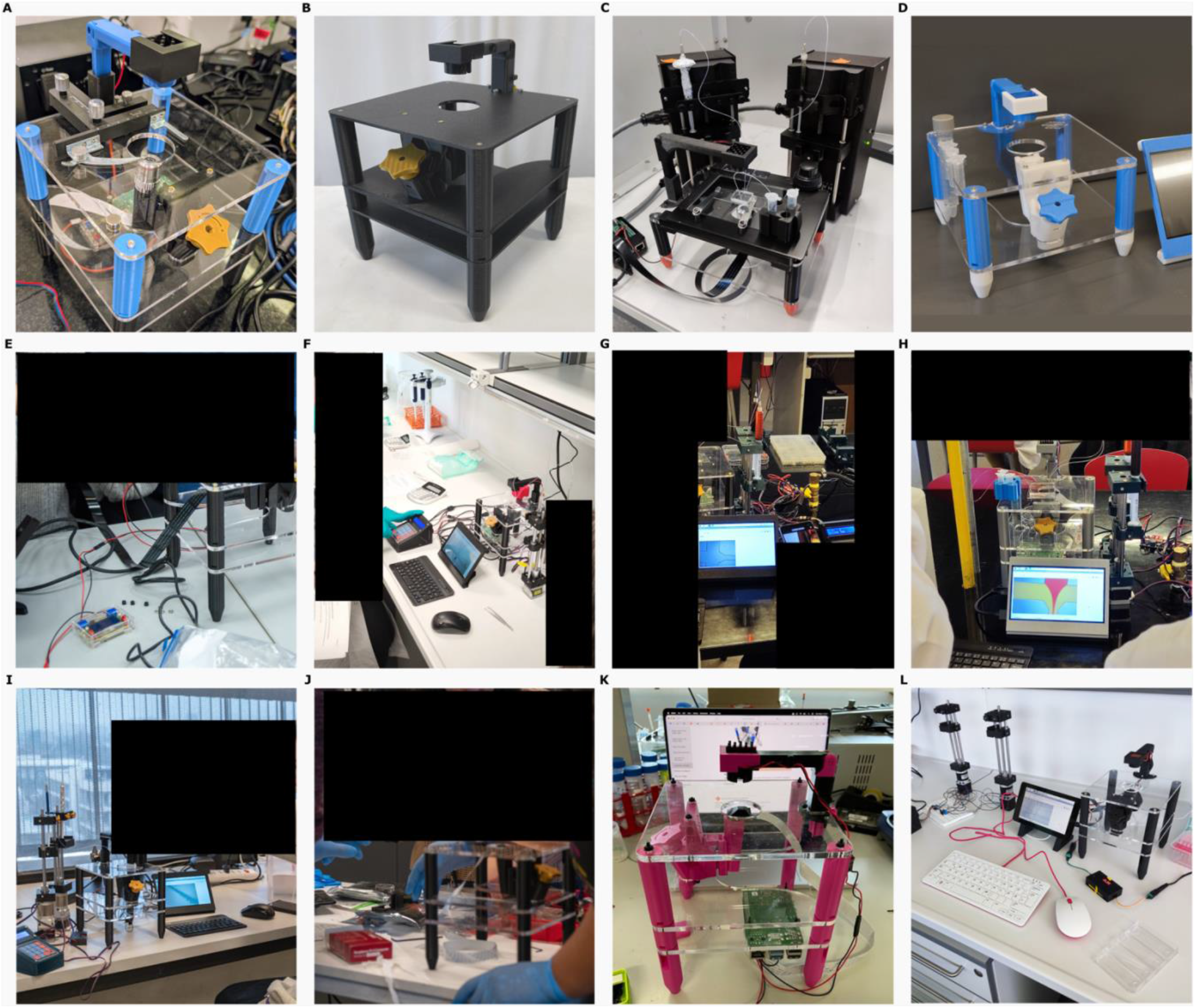
Design variants, training uses, and independent replication of the open-source droplet microfluidic workstation. (A) Standard three-level acrylic stage with a manual XY slide holder. (B) Fully 3D-printed alternative. (C) Adaptation for platform XY translation. (D) Compact two-level design with integrated tube holders. (E) Assembly during one of several LIBRE Hub workshops. (F) Experimental training using our platforms at EMBO Practical Courses *Microbial metagenomics: a 360° approach*. (G,H) Undergraduate teaching of droplet generation and diffusion in laminar-flow. (I,J) two of six Latin American senior researchers from Chile, Argentina and Mexico finishing their new instruments in LIBRE hub workshops. (K) Largely independent replication in the Netherlands, Saggiomo Lab. (L) Fully independent build at the EMBL Heidelberg, including electronics and pumps, from our released documentation.

The microfluidic platform has been used for professional and academic teaching instrumentation building (**Fig. 8E**), professional training of advanced research methods at three EMBO courses (**Fig. 8F**), and hands-on droplet microfluidic applications in university classes (**Fig. 8G, H**). All of these topics are usually out of reach for group hands-on teaching due to the instrumentation requirements.

To evaluate the replicability of the platform, intra- and inter-laboratory rebuilds were assessed through hands-on workshops and user reports (**Fig. 8E, I, J**). Within the Latin American Hub for Bioimaging through Open Hardware (LIBRE Hub), hands-on workshops held in Santiago, Chile, brought together graduate students and senior researchers from Chile, Argentina, and Mexico to independently assemble the microscopy stage using only the publicly available documentation and bills of materials. Across two workshops, six fully operational strobe microscopy systems were constructed, demonstrating that researchers without prior involvement in the hardware development could reproduce the instrument using the released documentation.

Replicability was further demonstrated by a partially independent fabrication and assembly of the microscopy stage in The Netherlands (**Fig. 8K**), and a fully independent reconstruction of the complete platform in Heidelberg, Germany, where both the microscopy stage and syringe-pump system were fabricated, assembled and used for research and training in custom single-cell genomics applications (**Fig. 8L**). Additional independent builds have been reported by external users, especially of the syringe-pump system through the Printables community. Although these examples represent a limited number of independent implementations, they demonstrate the adoption beyond the originating laboratory.

Instead of concluding this open-source hardware study with the public release of design files, the present work provides evidence that independent users can successfully reconstruct and operate the proposed instrumentation across different laboratories and geographical regions. Combined with the performance characterization, automation capabilities, and biological validation presented throughout this study, these replication efforts support the broader objective of establishing accessible, reproducible, and extensible open-source infrastructure for droplet microfluidics and single-cell research.

## Conclusions

This study presented the design, fabrication, and validation of an accessible open-source droplet microfluidics platform integrating a strobe-enhanced microscopy stage with digitally fabricated syringe pumps. Across the operating conditions examined, the optimised platform achieved flow accuracy, stability, and droplet monodispersity comparable to the commercial reference systems. In particular, the gearbox-assisted pump approached the performance of the Harvard Apparatus PHD 2000 and maintained stable delivery at low flow rates under high hydraulic resistance. Our microscopy stage performs all key microfluidics tasks without the need for a costly high-speed camera.

The modular architecture provides practical advantages beyond cost reduction. A shared web interface and API permit independent or coordinated control of the microscopy stage and pumps, while the pump keypad retains immediate manual control. Laser-cut and 3D-printed structures, modular electronics, replaceable motion components, and openly released design files enable local fabrication, repair, and adaptation. Software-based hardware simulation and detailed assembly documentation further reduce the technical requirements for implementation and adaptation, as demonstrated by successful builds independently completed outside the originating laboratory. Programmatic adjustment of multiple phase flow rates enabled automated flow sequences and controlled changes in droplet size and composition during continuous operation.

Biological validation demonstrated the platform’s ability to encapsulate and monitor individual cells and multicellular structures within microfluidic environments. The simple imaging system provided sufficient spatial and temporal resolution to assess droplet occupancy, bacterial growth dynamics, and basic cellular morphology over extended time-lapse experiments.

Together, these results establish the platform as a reliable and adaptable alternative to proprietary droplet microfluidics instrumentation. By combining validated experimental performance with open documentation, local manufacturability, repairability, and programmable control, the platform lowers technical and financial barriers to droplet-based research and provides a foundation for further automation and application-specific development.

## Materials and methods

### Microfluidic chip fabrication

Single-layer polydimethylsiloxane (PDMS) microfluidic chips were fabricated using conventional photolithography and soft lithography techniques, as previously reported^44^. Master molds were fabricated using photolithography on 100 mm diameter silicon wafers (Silicon Materials). Using soft lithography, single-layer replicas were produced by pouring a degassed PDMS and curing agent mix (10:1 ratio; Sylgard 184) onto the master molds and curing it at 80 °C for four hours. The PDMS replicas were carefully peeled off and punched at the inlet and outlet ports using a 0.75 mm biopsy punch. The replicas were then bonded to microscopy glass slides (Paul Marienfeld GmbH & Co. KG) after surface activation in an oxygen plasma oven (Diener Electronic GmbH) and coated with trichloro(1H,1H,2H,2H-perfluorooctyl)silane (Sigma) in fluorinated oil HFE-7500 (1% w/w; 3M) to render the microchannels hydrophobic. Once filled, the chips were heated at 75 °C for 20 min on a hotplate to evaporate the supernatant^45^.

The chip design was a droplet generation chip that used a flow-focusing junction in which the aqueous phase inlet was hydrodynamically focused by two lateral oil phase inlets converging at a nozzle. This design exhibited rectangular cross-section channels with a height of 40 μm, inlet channels of 105 μm width converging into a main channel of 245 μm width, with a total length of 15 mm approximately.

### Multi fluid phase manipulation

The flow accuracy and stability of the open-source syringe pump designs were systematically evaluated and benchmarked against a commercial laboratory syringe pump (PHD2000; Harvard Apparatus). Sensirion LG16-0431D flow sensors were integrated between the syringe pumps and microfluidic network to enable real-time flow monitoring. Custom Python scripts were developed for data acquisition and analyses. The open-source designs were operated using the same microstepping configuration (32 microsteps), whereas the Harvard Apparatus pump operated under the manufacturer’s default settings. Fluidic characterization was performed using 3 mL plastic syringes (Nipro Corporation; nominal inner diameter: 8.66 mm) with rubber plunger seals, distilled water, and polytetrafluoroethylene (PTFE) tubing (Adtech Polymer Engineering Ltd.; OD: 1/16”, ID: 0.75 mm).

To evaluate the pump performance under realistic experimental conditions, the droplet generation chips were tested at target flow rates of 25, 100, 500, and 1000 μL h^−1^. For each flow rate setpoint, 16000 data points (∼10 min) were acquired at a sampling rate of 27 Hz (sufficient to resolve pulsation frequencies up to 13.5 Hz per Nyquist criterion) following a 5-minute flow stabilization period to ensure steady-state conditions. When pulsations exceeded sensor bandwidth, they were assessed indirectly through baseline stability.

To assess multiphase flow control, proprietary and open-source syringe pump models and a droplet generation chip were used to produce water-in-oil droplets. Distilled water served as the discrete phase, and fluorinated oil (3M Novec 7500) with 1% PicoSurf surfactant (5% w/w in Novec 7500; Sphere Bio Ltd.) served as the continuous phase. Both fluids were loaded into 3 mL plastic syringes and connected to the ports of the microfluidic chip using PTFE tubing. By maintaining a 2:1 oil-to-aqueous flow rate ratio, droplet monodispersity was evaluated after a 2 min stabilization period at flow rates of 50:25, 200:100, 1000:500, and 2000:1000 μL h^−1^ using image-based analysis with automated boundary detection without manual selection (**Fig. 4**).

Dynamic modulation assays were performed using the high-torque pump that controlled the ratio of discrete and continuous phases. A red food dye and distilled water mixture (1:20 v/v ratio; Gourmet) was introduced as the second discrete phase before droplet formation. Two experiments were performed using this configuration: droplet size modulation and dilution modulation. For droplet size modulation, different droplet sizes were produced by automatically modulating the oil phase to 1000, 1500, and 2500 μL h^−1^, while maintaining the water-dye ratio (250:250 μL h^−1^). For dilution modulation, a gradient of droplet compositions was generated by progressively changing the water-red dye ratio through nine steps from 990:10 μL h^−1^ to 10:990 μL h^−1^ (red dye flow rates of 10, 125, 250, 375, 500, 625, 750, 875, and 990 μL h^−1^), while maintaining the oil flow rate at 1500 μL h^−1^.

### Bacterial and microsphere encapsulation

Following droplet generation, the platform was used to encapsulate bacteria and calibration microspheres within water-in-oil droplets and to monitor their behavior within these microenvironments. *Escherichia coli* TOP10 and *Staphylococcus aureus* strains were selected as the primary biological models for bacterial encapsulation. These strains harbored plasmids encoding a superfolder green fluorescent protein (sfGFP) and kanamycin resistance^27^ , and GFP and tetracycline resistance, respectively.

To prepare the cell suspensions for encapsulation, bacterial strains were streaked from frozen stocks onto agar plates prepared with pre-mixed Luria-Bertani Broth Miller (LB; United States Biological) supplemented with kanamycin (100 µg/mL) for sfGFP-expressing *E. coli*, and pre-mixed BBL Tryptic Soy Broth (TSB; Becton, Dickinson and Company) supplemented with tetracycline (12.5 µg/mL) for GFP-expressing *S. aureus*. Plates were incubated overnight at 37 °C. Individual colonies were picked using a sterile loop and used to inoculate 5 mL of liquid LB or TSB containing the corresponding antibiotic in separate culture tubes. Following overnight incubation at 37 °C, 50 µL of each culture was transferred into a new sterile tube containing 5 mL of fresh medium with the appropriate antibiotic to re-establish exponential growth. Cultures were incubated at 37 °C with shaking at 250 rpm until reaching an optical density at 600 nm (OD600) of 0.5, measured using a Jenway 6320D spectrophotometer.

Prior to microfluidic experiments, sfGFP-expressing *E. coli* and GFP-expressing *S. aureus* cultures were diluted to a final concentration of 1.41e+7 cells/mL. Cell concentrations were calculated assuming Poisson encapsulation statistics, targeting an average occupancy of one cell per five droplets and a droplet volume of 14.14 pL (30 µm diameter droplets)^27^. For microsphere encapsulation experiments, 20 µL of 10.5-µm Ultra Rainbow calibration particles (Spherotech, Inc.) were diluted in 1 mL of NaCl solution. The suspension was vortexed for 1 min to ensure homogeneous particle dispersion, resulting in a final concentration of approximately 1.96 e+5 particles/mL.

Bacterial and microsphere suspensions were loaded into 1 mL plastic syringes (BH Supplies), while the oil phase was loaded into 3 mL plastic syringes. Both fluid phases were connected to the microfluidic chip inlets using PTFE tubing. The outlet ports were similarly connected to either waste containers or collection tubes (2 mL Eppendorf tubes). A 2:1 oil-to-aqueous flow rate ratio was used to generate approximately 30 µm diameter droplets within the dripping regime of the flow-focusing geometry.

### Imaging and analysis

Bright-field microscopy was performed using the strobe-enhanced microscopy stage and 4 ×, 20 ×, and 40 × objectives (tube length: 160 mm / coverglass: 0.17 mm). Sequential on-chip images were acquired during droplet generation, whereas off-chip images were obtained for encapsulated samples. Glass observation chambers were assembled to image the resulting emulsions and monitor dynamic events^46^. For single-timepoint imaging, droplet monolayers were formed by sequentially loading 5 μL of Novec oil, 3 μL of emulsion, and an additional 15 μL of oil into the observation chamber to facilitate droplet distribution. When time-lapse microscopy was required, microdroplet monolayers were loaded, and a sealant was applied to the chamber to close any openings and prevent evaporation of the emulsion during extended imaging experiments.

To analyze droplet populations, the raw frames stacked into datasets were processed using a custom Python-based computer vision tool (https://github.com/wenzel-lab/droplet_AInalysis) to detect droplet boundaries and define these areas as regions of interest (ROI) for further morphology (e.g., size and circularity) and composition (e.g., intensity) analysis. In dilution experiments, mean green-channel intensity was quantified as a function of the food-dye-to-water flow-rate ratio to assess optical absorbance. For biological assays, 12-h time-lapse image sequences were acquired to monitor the growth of sfGFP-expressing *E. coli* and GFP-expressing *S. aureus* within droplets. Experiments were performed at room temperature (18-22 °C), enabling continuous long-term imaging without a heated environmental enclosure.

All microscopy images presented in this study were acquired using the Raspberry Pi Camera V2 with stroboscopic illumination, as described above.

## Author contributions

PPH and TW developed the methodology, performed the investigation, curated and analysed the data, validated the results, prepared the visualisations, administered and supervised the project, and drafted the manuscript. JBRA contributed data. TW conceived the study, and acquired funding. PPH, CN, TA, WA, JJL and TW developed the software. All authors provided design resources, reviewed and edited the manuscript.

## Data availability

Data for this article, including the design files and assembly instructions are available on GitHub:

Strobe-enhanced microscopy stage https://github.com/wenzel-lab/strobe-enhanced-microscopy-stage, 3D-printed syringe pumps https://github.com/wenzel-lab/syringe-pumps-and-controller/, and controller boards and software https://github.com/wenzel-lab/rio-controller).

Data and notebooks from the characterization and analysis are available on the Open Science Framework: https://osf.io/mjyab.

## Acknowledgements

This work was supported by the Chan Zuckerberg Initiative funding to the LIBRE hub project. This research was furthermore supported by ANID FONDECYT Regular 1241621, and Centros de Investigación y Desarrollo de Excelencia de Interés Nacional, ANID SENTINET CIN250062. We also thank all researchers who replicated our designs internationally who reached out to share images and experiences.

