## Supplementary figures for "An open-source, 3D-printable droplet microfluidics platform with strobe-enhanced microscopy"

[a] Institute for Biological and Medical Engineering, Schools of Engineering, Medicine and Biological Sciences, Pontificia Universidad Católica de Chile,Santiago, Chile
[b] Chipbotics, London, United Kingdom.
[c] IOWlabs, Santiago, Chile

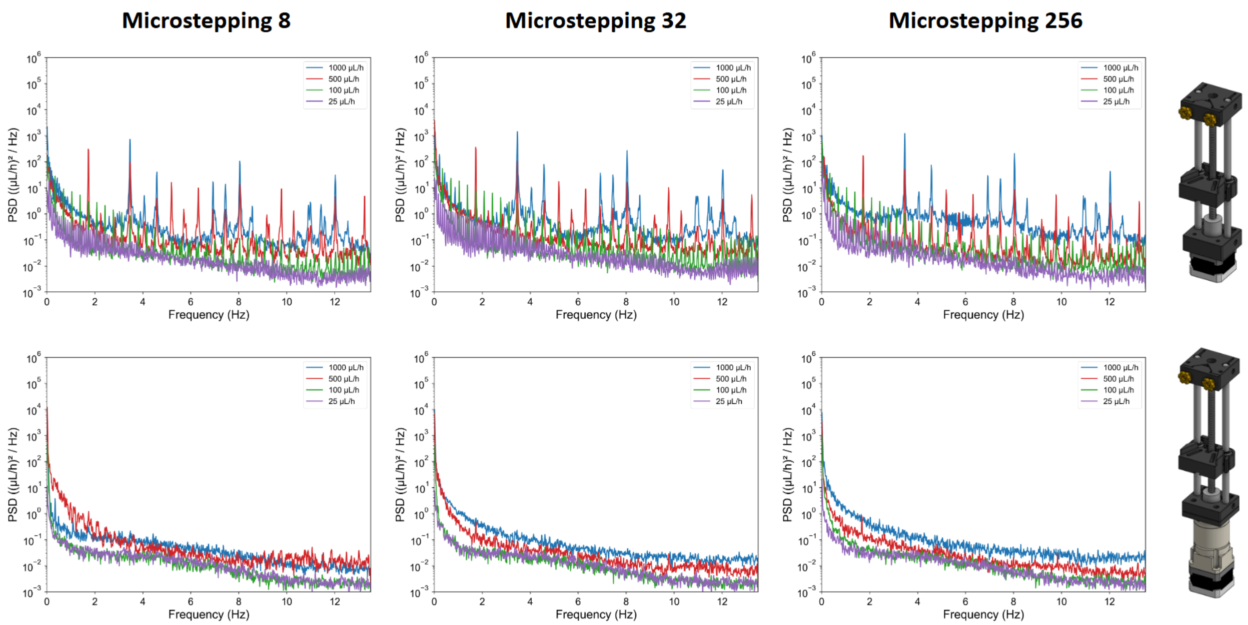


**Figure S1 |** Power spectral density (PSD) analysis of flow rate fluctuations for the two open-source syringe pump architectures operated at different microstepping resolutions (8, 32, and 256 microsteps) and target flow rates (25, 100, 500, and 1000 µL $h^{-1}$).

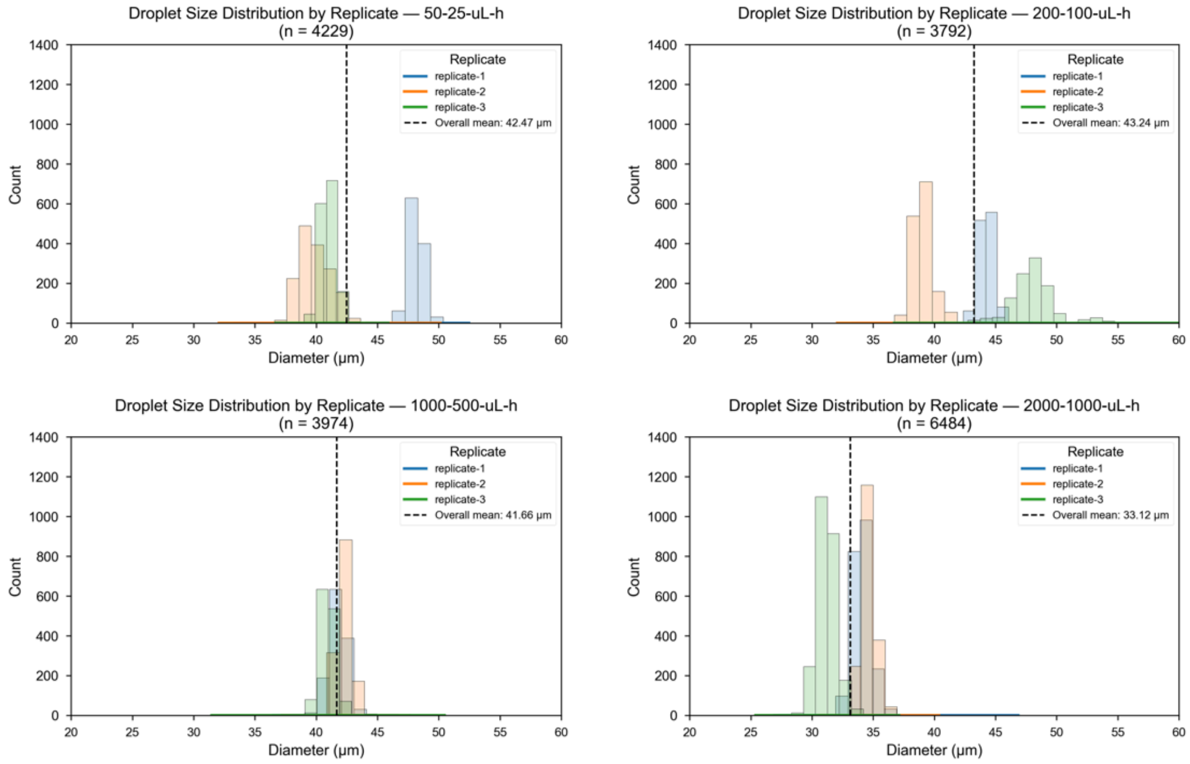


**Figure S2 |** Droplet size distributions obtained from three technical replicates using the gearbox-assisted syringe pump at four representative oil-to-aqueous flow-rate combinations (50:25, 200:100, 1000:500, and 2000:1000 µL $h^{-1}$).

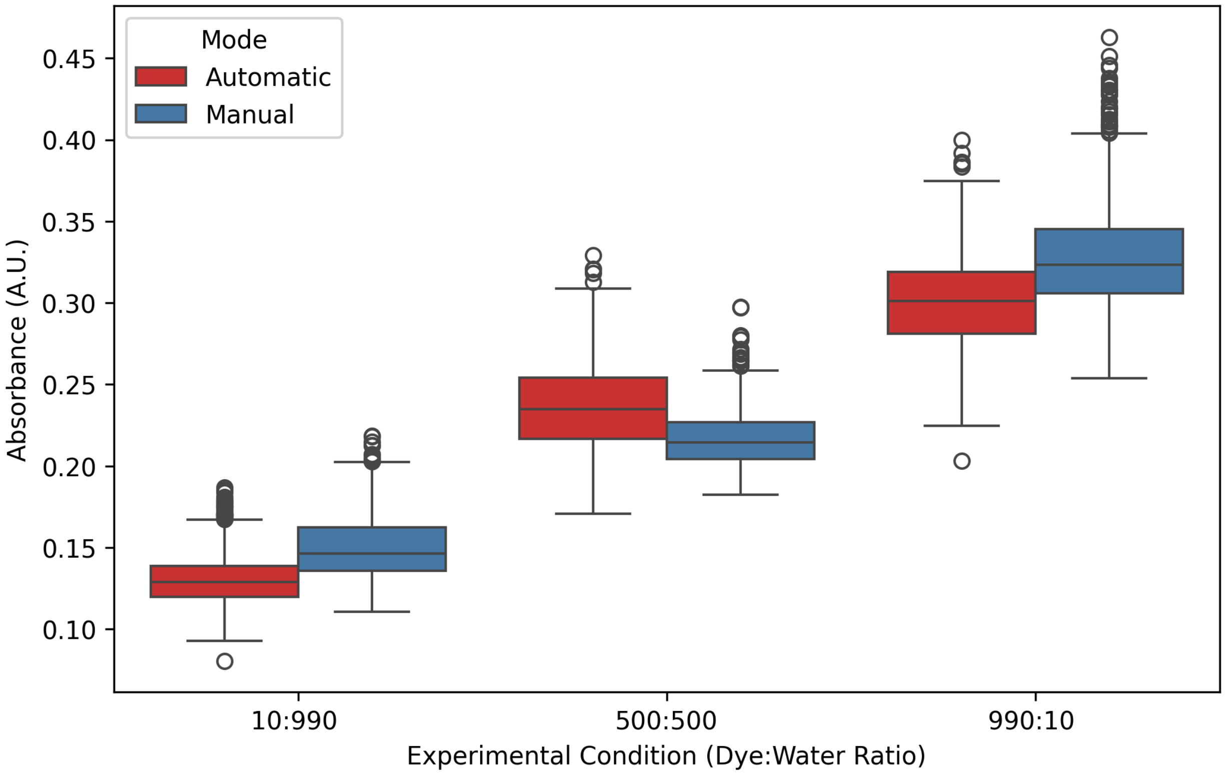


**Figure S3 |** Comparison of dilution modulation assays performed using the gearbox-assisted syringe pump in automatic and manual operating modes.
